# A Spray Dried Replicon Vaccine Platform for Pandemic Response

**DOI:** 10.1101/2025.04.10.648241

**Authors:** Wynton D. McClary, John Z. Chen, Hui Wang, Ethan Lo, Julie Bakken, Joseph McCollum, Christopher Press, Eduard Melief, Devin S. Brandt, Andrew R. Martin, Reinhard Vehring, Darshan N. Kasal, Emily A. Voigt, Alana Gerhardt

**Author notes:** Corresponding Author: Alana Gerhardt.

## Abstract

The recent COVID-19 pandemic, as well as the threat of a global pandemic caused by H5N1 avian influenza virus, has highlighted the need for the development of thermostable vaccines that can be manufactured and distributed rapidly to combat the next global pandemic. To address this need, we previously developed a replicon vaccine platform that utilizes a nanostructured lipid carrier (NLC) to protect and efficiently deliver antigen-expressing replicon molecules *in vivo*. The replicon-NLC vaccine platform uses readily sourced components and can be rapidly manufactured at scale with the potential for stockpiling, thus enhancing pandemic preparedness. Spray drying is a promising method of vaccine desiccation with reduced costs and increased scale-up capabilities compared to lyophilization. As proof of concept, we demonstrate for the first time the successful spray drying of a replicon-NLC vaccine complex designed to protect against H5N1 avian influenza A virus to enhance its long-term thermostability while maintaining vaccine immunogenicity in an *in vivo* mouse model. Several glass-forming disaccharide excipients were screened for formulation and process compatibility under low-temperature spray drying conditions, and it was determined that a suitable shell-forming excipient, L-leucine, was necessary to prevent excessive accumulation of replicon-NLC vaccine complexes on the dry powder surface and a subsequent loss in process yield. The spray dried replicon-NLC vaccine powders were chemically stable for 1 month of storage at 40°C. Immunogenicity of the spray dried drug product was also well maintained for at least 3 months of storage at 4°C when administered intramuscularly into C57BL/6 mice as a reconstituted liquid. Finally, we demonstrate the ability to precisely control the aerodynamic particle size of the spray dried vaccine product to generate dry powders that are theoretically suitable for nasal or pulmonary delivery without reconstitution. This work establishes the feasibility of spray drying a thermostable replicon-NLC vaccine for rapid pandemic response.

## INTRODUCTION

Pandemics remain an ever-present threat to global health and safety. The COVID-19 pandemic, in particular, has highlighted key areas of pandemic response that are currently lacking, notably, the lack of cold-chain maintenance in rural or remote locations, both within the U.S. as well as in developing countries [1–3]. Further complicating matters, most prophylactic vaccines are administered via intramuscular injection, which requires trained medical staff to properly administer. However, due to a global shortage of health workers that is further exacerbated by inequities in the global distribution of this workforce, many developing and low-income countries simply lack enough trained personnel to properly administer vaccines efficiently in the event of a pandemic [4]. In addition, needle anxiety is common among the global populace, and needle-free methods of vaccine administration, such as intranasal administration of dry powder vaccines, will be essential to maximize the effectiveness of future vaccination campaigns [5,6]. With new and emerging pandemic threats, such as the avian influenza A H5N1 virus that has been spreading through domestic and wild animal populations and is beginning to cause human infections [7–13], it is more important than ever that new strategies are developed to address the common global distribution challenges that are associated with many vaccines.

Nucleic acid vaccines have shown exceptional utility during the COVID-19 pandemic as a tool for pandemic response, enabling rapid adaptability to new pathogen targets. In particular, the use of RNA vaccine technology eliminates the need for tedious and resource-limited vaccine production in embryonated chicken eggs [14], the standard production technique for current seasonal influenza vaccines. However, current RNA vaccine technologies have several key limitations that need to be addressed to enable complete global accessibility in the event of future pandemics. First, RNA vaccines are currently administered through intramuscular injection, which induces both humoral and cellular immunity that protect against viremia and serious disease but not against asymptomatic infection and viral shedding at the primary site of infection [15,16]. Second, the majority of currently approved RNA vaccines use lipid nanoparticles (LNPs) as the delivery vector, which often require complex manufacturing processes, difficult-to-source ionizable lipids, and encapsulation of the RNA during LNP manufacture [17] – complicating technology transfer and supply-chain management. Finally, currently marketed RNA vaccines require cold-chain storage (−20°C or −80°C), significantly impacting vaccine distribution to low-resource locations in particular [18,19].

To address these limitations, we have developed a novel nanostructured lipid carrier (NLC)-based replicon vaccine delivery system that uses readily-sourced lipids and surfactants, and can be quickly manufactured in a scalable manner using standard processes and equipment [20–23]. We have previously used this platform to develop an intramuscular vaccine against SARS-CoV-2 that successfully completed Phase 1 clinical trials [24]. However, due to the thermolabile nature of RNA in general, frozen storage is necessary to maintain the long-term stability of the replicon-NLC complex and other liquid RNA vaccine products. For this reason, various drying techniques have seen increased use in vaccine development to enable long-term storage of particularly sensitive formulations at ambient temperatures [25]. For example, we previously demonstrated that lyophilization of the replicon-NLC platform greatly enhances its thermostability over the course of several months [22]. Elsewhere, Zimmerman et al. demonstrated the successful spray drying of RNA-containing LNPs, which remained stable over the course of 3 months at ambient temperatures [26]. In the context of vaccine administration, a spray dried replicon-NLC vaccine powder would be amenable to a needle-free approach using intranasal or pulmonary routes of delivery to the local mucosa [27,28]. This would enable vaccination at the primary site of infection, a strategy that has previously been associated with decreased viral transmission for respiratory pathogens [29–32], and would have the added benefit of increasing vaccine uptake among the global population due to the reduced need for trained clinicians and needle-based administration strategies in more remote locations. There are several examples in the literature for the successful spray drying of RNA. For instance, Friis and co-workers were able to spray dry mRNA-containing LNPs and demonstrated intratracheal delivery using an insufflator for delivery [33]. Sarode and co-workers were separately able to achieve a similar result by spray drying a different mRNA-containing LNP formulation [34]. Elsewhere, Qiu and co-workers demonstrated successful spray drying and pulmonary delivery of mRNA using a PEGylated synthetic peptide as a delivery vector [35]. However, to our knowledge, there are no examples of a functional spray dried replicon vaccine with the potential for intranasal delivery.

Here, we establish proof-of-concept spray drying of a replicon-NLC vaccine using an H5 influenza antigen-expressing replicon as a model antigen. Several formulation conditions were screened to identify key excipients necessary to achieve a high-quality spray dried powder with a suitable process yield, and the physicochemical and immunogenic properties of the spray dried vaccine were compared to the liquid vaccine feedstock control. We next tested the thermostability of the dry powder vaccine presentation relative to the liquid presentation after several months of storage at various temperatures. Finally, we examined whether the size of the manufactured dry powder could be controlled to enable direct intranasal or pulmonary administration of the spray dried vaccine.

## MATERIALS AND METHODS

### Manufacturing of NLCs

The NLC formulation was prepared according to an established protocol for research-grade NLC [20–22,36–39]. Briefly, squalene (MilliporeSigma, Burlington, MA, USA), sorbitan monostearate (Span 60; Spectrum Chemicals, New Brunswick NJ, USA), 1,2-dioleoyl-3-trimethylammoniumpropane (DOTAP; SAFC, St. Louis, MO, USA), and trimyristin (Dynasan 114; MilliporeSigma) were mixed and dissolved by heating to 65°C and sonication using a water bath sonicator to form the oil phase. Separately, polysorbate 80 (Tween 80; MilliporeSigma) was mixed and dissolved in a solution of 10 mM sodium citrate by heating to 65°C with water bath sonication to form the aqueous phase. After both the oil phase and aqueous phase were fully dissolved, the two were rapidly mixed using a high-speed laboratory emulsifier (Silverson Machines, Chesham, UK), and 10 discrete microfluidization cycles were carried out on the resultant suspension using a microfluidizer (M-110P, Microfluidics International Corp., Westwood, MA, USA). The final NLC product contained 30 mg/mL DOTAP, 37.5 mg/mL squalene, 2.4 mg/mL trimyristin, 37 mg/mL polysorbate 80, and 37 mg/mL sorbitan monostearate. NLCs were terminally filtered using a 0.22 µm polyethersulfone filter, and the Z-average hydrodynamic diameter was confirmed to be 60 ± 25 nm by dynamic light scattering (DLS; described below). NLCs were stored at 2 – 8°C in 3 mL glass vials until further use.

### Replicon synthesis and purification

H5 replicon synthesis and purification was performed using a previously established method [21–23,36–40] with minor modifications. Briefly, replicon template DNA plasmids were generated to contain a codon-optimized influenza A/Vietnam/1203/2004 H5N1 hemagglutinin (HA) antigen (GenBank EU122404.1), as well as the 5’ UTR, 3’ UTR, and non-structural proteins (nsp1-4) derived from the attenuated TC83 strain of Venezuelan equine encephalitis virus. Template DNA was linearized by NotI restriction digestion for *in vitro* transcription (IVT) of the replicon. Upon completion of IVT, DNase I (Aldevron, Fargo, ND, USA) was used to digest the remaining template DNA plasmid, and cap0 structures were added to the RNA replicon transcripts using vaccinia capping enzyme (Aldevron). The capped replicon was purified using Capto Core 700 multimodal chromatography resin (Cytiva, Marlborough, MA, USA), followed by diafiltration and concentration using tangential flow filtration. The final replicon material was terminally filtered using a 0.22 µm polyethersulfone filter. Purity was assessed by agarose gel electrophoresis, and the replicon was quantified via a NanoDrop (Thermo Fisher Scientific, Waltham, MA, USA) as well as a commercially available Quant-it RiboGreen RNA assay (Thermo Fisher Scientific). The replicon was stored at −80°C until further use.

### Formation of H5 hemagglutinin replicon vaccine complex with NLCs

H5 replicon-NLC vaccine complexes were formed as described previously [21,22], with minor modifications. All vaccine complexes were prepared using a nitrogen:phosphate (N:P) ratio of 10 (N:P represents the ratio of cationic quaternary ammonium ions of the DOTAP lipid component of the NLC to the anionic phosphate groups of the nucleic acid backbone). Diluted replicon was mixed 1:1 by volume with diluted NLC to achieve vaccine complexes at the desired replicon nucleic acid concentration in a buffer containing 10 mM sodium citrate in the presence of either 10% w/v trehalose, 9% w/v lactose, or an 8%/2% trehalose/sucrose mixture. For formulations containing L-leucine, all formulation components (replicon, NLC, 10 mM sodium citrate, and 10% w/v trehalose) were prepared in a solution of 20 mg/mL L-leucine, dissolved in nuclease-free water (Invitrogen, Waltham, MA, USA). All replicon-NLC complexes were allowed to incubate on ice for 30 minutes prior to further use.

### Spray drying of replicon-NLC complexes

Spray drying of vaccine complex was carried out using a small-scale spray dryer (4M8-Trix PROCEPT Particle Processing Equipment, Zele, Belgium). To expedite the selection of suitable spray drying process conditions, a theoretical process model for the spray dryer was utilized, which takes into account the dimensions and heat loss of the spray dryer, the liquid feedstock and drying gas flow rates, the feedstock solids concentration, the atomizing gas flow rate and pressure (if applicable), the drying gas inlet temperature, the drying gas relative humidity to predict the dryer outlet temperature and air humidity, and the moisture content of the spray dried powder [41]. Process parameters for dry powders produced using the twin fluid atomizer are summarized in **Table 1**. To achieve a dry powder aerodynamic particle size distribution suitable for pulmonary delivery (< 5 µm), the spray dryer was equipped with a twin fluid atomizer with an orifice diameter of 0.4 mm (PROCEPT Particle Processing Equipment). To minimize the potential for thermal-induced degradation of the replicon and NLC components during the drying process, process parameters were chosen to achieve a low outlet temperature of approximately 43°C and a moisture content below 4%. The inlet temperature was set to 65°C, the drying gas flow rate was set to 400 standard liters per minute (SLPM), and the inlet drying gas relative humidity (RH) was measured to be 15%. The atomizing gas flow rate was set to 10 L/min (corresponding to an atomizing gas pressure of approximately 2.25 bar) using compressed dry air as the atomizing gas, and the feedstock was atomized at a liquid flow rate of 1 mL/min. This resulted in an air-to-liquid ratio of 11.8 and an initial median droplet diameter of approximately 7.3 µm (span = 1.4), as measured by a Spraytec droplet size measurement system (Malvern Panalytical, Worchestershire, UK). The selected spray drying process parameters corresponded to a predicted outlet gas RH of 9.1% and a predicted residual moisture content of 3.1%. After spray drying, the powder was transferred to a clean and dry glass scintillation vial and stored in a dry box at less than or equal to 14% RH short-term or aliquoted into clean glass scintillation vials and stored in double heat-sealed aluminum foil pouches with molecular sieve desiccant for longer-term storage at varying temperatures. The recovered powder yield, *E*, was calculated according to the following equation:

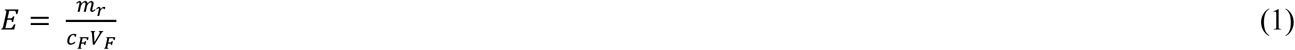

where *m*_r_ is the recovered powder mass and *c*_F_ is the total solids concentration of the feedstock with volume *V*_F_. The predicted mass median aerodynamic diameter (MMAD) *d*_a,50_ of the powder was calculated according to the following equation [42–44]:

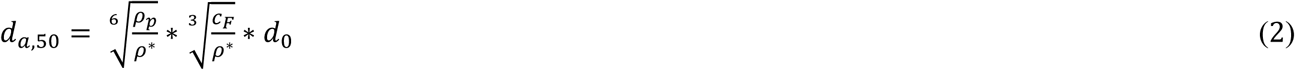

where *ρ*_p_ is the particle density, *ρ*\* is the unit density (1000 g/L), *c*_F_ is the feedstock concentration, and *d*_0_ is the initial droplet diameter. Using laser diffraction analysis to determine the median volume equivalent diameter (*d*_v,50_) of the dry powder (described below), the particle density *ρ*_p_ was calculated according to the following equation:

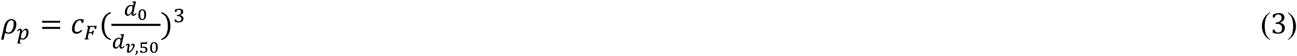

**Table 1.**
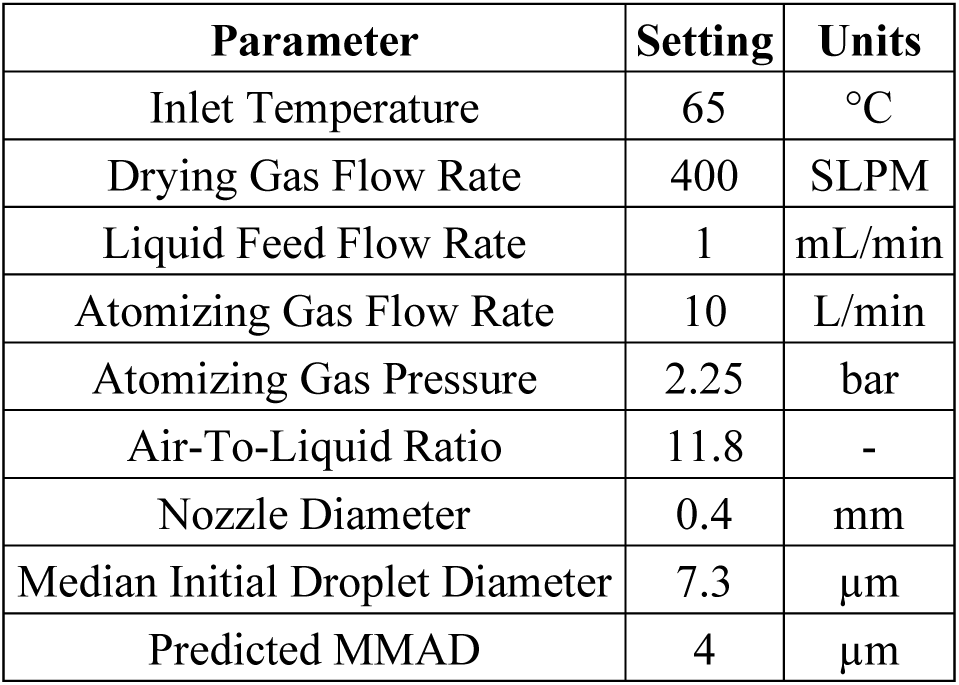
Twin fluid atomizer spray drying process parameters and predicted MMAD.

| Parameter | Setting | Units |
| --- | --- | --- |
| Inlet Temperature | 65 | °C |
| Drying Gas Flow Rate | 400 | SLPM |
| Liquid Feed Flow Rate | 1 | mL/min |
| Atomizing Gas Flow Rate | 10 | L/min |
| Atomizing Gas Pressure | 2.25 | bar |
| Air-To-Liquid Ratio | 11.8 | - |
| Nozzle Diameter | 0.4 | mm |
| Median Initial Droplet Diameter | 7.3 | µm |
| Predicted MMAD | 4 | µm |

To achieve a dry powder particle size greater than 20 µm (suitable for intranasal delivery), the spray dryer was equipped with a 25 kHz ultrasonic atomizer to generate a larger initial droplet diameter of approximately 65 µm (**Table 2**). To increase the residence time of the droplets in the chamber to allow for sufficient drying, a low drying gas flow rate of 250 SLPM was used. An additional 150 SLPM of cool, dry air (yielding a total cyclone separator air flow rate of 400 SLPM) was applied at the U-bend to lower the humidity at the collector and adjust the cyclone cutoff to boost powder recovery. The liquid feed flow rate and inlet temperature were kept the same as for the material spray dried using the twin fluid atomizer. The recovered powder yield and predicted MMAD were calculated according to Eqs. 1, 2, and 3 as described above.

**Table 2.** Ultrasonic atomizer spray drying process parameters and predicted MMAD.

| Parameter | Setting | Units |
| --- | --- | --- |
| Inlet Temperature | 65 | °C |
| Drying Gas Flow Rate | 250 | SLPM |
| Secondary Gas Flow Rate | 150 | SLPM |
| Total Flow Rate in Cyclone | 400 | SLPM |
| Liquid Feed Flow Rate | 1 | mL/min |
| Ultrasonic Frequency | 25 | kHz |
| Initial Droplet Diameter | 65 | μm |
| Predicted MMAD | 35 | μm |

### Laser diffraction

To determine the *d*_v,50_, optical size distributions of the primary particles were determined via wet laser diffraction (Partica LA-960V2, Horiba, Ltd., Kyoto, Japan), which incorporates the full Mie scattering theory. 1-octanol (99% 1-octanol; Thermo Fisher Scientific; refractive index *n* = 1.429 - 0.000*i*) was used as a non-solvent liquid dispersion medium. Measurements were conducted as follows: first, approximately 200 mL of 1-octanol was loaded into the clean measurement cell. Spray dried powder (refractive index *n* = 1.652 – 0.000*i* [45]) was then added incrementally until the laser transmittance fell below 90% and the residual parameter value *R* fell below 0.10, according to instrument manufacturer recommendations. A 60-second sonication step was then applied to the dispersion medium to de-agglomerate particle aggregates, followed by a 10-minute rest step to allow the remaining bubbles to dissipate.

### Aerodynamic particle size measurements

The aerodynamic particle size of the spray dried particles was measured using an Aerodynamic Particle Sizer (APS) (3321, TSI, Shoreview, MN, USA) equipped with a small-scale disperser (3433, TSI). Powders were loaded onto annular rings of abrasive paper glued to the top of a turntable and brushed gently for improved dispersion. For all measurements, the APS was operated for a total of 180 s for each measurement. All samples were measured in triplicate, and data were averaged.

### Karl Fischer titration

The moisture content of spray dried powders was determined using a coulometric titrator equipped with an oven attachment (C30SD, Mettler Toledo, Columbus, OH, USA). Glass sample vials were filled with approximately 100 mg of powder, sealed with plastic screw caps, and placed into the oven attachment. The instrument was corrected for drift using an empty vial, and three empty vials were assessed for blank subtraction prior to sample analysis. Samples and blanks were heated to 130°C, and water release was analyzed for a maximum of 20 min to determine the moisture content in units of percent total mass of the material.

### Scanning electron microscopy

High-resolution micrographs of the spray dried particles were obtained using a field emission scanning electron microscope (FESEM; Sigma, Zeiss, Germany) to examine particle morphology, with some particles intentionally cracked to reveal internal structures. A scanning electron microscope (SEM; JSM-IT100, JEOL, Peabody, MA, USA) was used to monitor general morphological changes during long-term stability studies. For all imaging, samples were mounted on standard SEM pin mounts (Ted Pella Inc., Redding, CA, USA) with double-sided adhesive carbon tape and sputter-coated with a thin layer of gold nanoparticles to eliminate charging effects. The coated samples were imaged in high vacuum mode using an acceleration voltage of 5 keV and a secondary electron detector.

### Raman spectroscopy

The solid phase of the individual components in the spray dried samples was analyzed using a custom-designed dispersive Raman spectrometer [46]. The instrument was equipped with a 671 nm diode laser (Ventus 671, Laser Quantum, Stockport, UK) with a maximum output power of 500 mW. Powder samples were loaded into a 0.2 µL conical cavity in an aluminum sample holder and exposed to the laser in a sample chamber flushed with clean, dry nitrogen (less than 3% RH) at room temperature. Raw crystalline L-leucine and crystalline trehalose dihydrate were measured directly to obtain their reference spectra. A spray dried batch of pure trehalose was used to generate the reference spectrum for amorphous trehalose. Since fully amorphous leucine could not be easily produced, its amorphous reference spectrum was obtained by subtracting the amorphous trehalose reference from the spectrum of a spray dried trehalose–leucine powder, with further details reported elsewhere [47].

### Dynamic vapor sorption analysis

To investigate the interaction between ambient moisture and the spray dried powder, which can affect the product’s long-term thermostability, a gravimetric water vapor sorption analyzer (DVS Intrinsic Plus, Surface Measurement Systems, Allentown, PA, USA) was used to measure the water sorption behavior of the powder product. Approximately 30 mg of powder was loosely dispersed onto the sample pan and equilibrated at 0% RH and 25°C to remove residual moisture. The moisture sorption and desorption isotherms were measured at 25°C, ranging from 0% RH to 90% RH, in 10% RH increments. At each stage, the equilibrium criterion was defined as a weight change of less than 0.002 %/min, after which the powder was considered to have equilibrated at the set RH, allowing the test to proceed to the next step.

### Dry powder reconstitution

Dry powder samples were reconstituted by the addition of room temperature nuclease-free water (Thermo Fisher Scientific). To account for the varying solids contents in the different tested feedstock formulations, the reconstitution volume *V*_S_ was calculated according to the following equation:

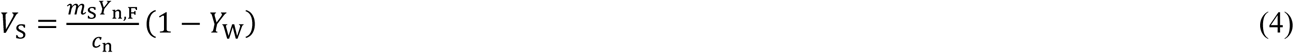

where *m*_S_ is the powder sample mass, *Y*_n,F_ is the mass fraction of nucleic acid as formulated in the feedstock, *c*_n_ is the target nucleic acid concentration in the reconstituted liquid, and *Y*_W_ is the mass fraction of water in the powder (relative to total powder mass). Using this equation, the appropriate volume of room temperature nuclease-free water was added to the spray dried powder to achieve the desired final nucleic acid concentration. Samples were vortexed briefly to resuspend the powder in solution (less than 10 sec) and allowed to incubate at room temperature until fully reconstituted to a liquid suspension (approximately 4 h).

### pH measurements

Sample pH measurements were carried out using a calibrated pH meter equipped with a ROSS micro-electrode probe (Thermo Fisher Scientific). Measurements were collected using 80 µL of neat liquid or reconstituted sample, and the probe was washed with oil and grease removal solution and rinsed with Milli-Q water in between each sample to prevent carryover.

### Dynamic light scattering and electrophoretic light scattering

Nanoparticle hydrodynamic diameter and zeta potential were determined by DLS and electrophoretic light scattering (ELS) in tandem using a Zetasizer Pro instrument equipped with a 633 nm laser (Malvern Panalytical). Liquid samples or dry powder samples (reconstituted to a replicon nucleic acid concentration of 50 µg/mL with Milli-Q water) were diluted 100-fold using nuclease-free water and loaded into a zeta cell (DTS1070 folded capillary zeta cell, Malvern Panalytical). Hydrodynamic diameter measurements were carried out at 25°C with a dispersant and sample viscosity value of 0.8872 cP (water). All measurements were collected using a backscatter angle of 173°. Three repeat measurements were collected and averaged, and the standard deviation was calculated.

Zeta potential measurements were carried out at 25°C with a viscosity value of 0.8872 cP, a refractive index of 1.330, and a dielectric constant of 78.5. Zeta potential data were collected using the Smoluchowski model [48] and a Henry function F(ka) value of 1.50. The measurement duration, attenuation selection, and voltage were each set to automatic, and a total of 5 repeat measurements were collected with a 60-second delay between repeat measurements. A minimum of 10 runs and a maximum of 30 runs were collected per repeat measurement. Repeat measurements were averaged, and the standard deviation of the mean was calculated.

### Agarose gel electrophoresis

Full-length replicon nucleic acid content in the liquid or reconstituted spray dried vaccine was assessed using an agarose gel-based method described previously [21]. Briefly, replicon-NLC complexes, or control H5 replicon alone (freshly thawed from −80°C storage), were diluted to a nucleic acid concentration of 40 µg/mL and subsequently diluted 1:1 by volume with 25:24:1 phenol:chloroform:isoamyl alcohol (Thermo Fisher Scientific). Samples were briefly mixed by vortexing and subsequently centrifuged at 17,000 *g* for 15 min to separate the organic and aqueous phases. The aqueous phase was extracted and mixed 1:1 by volume with glyoxal load dye (Invitrogen), heated at 50°C for 20 min, then allowed to cool to room temperature for 5 min prior to loading on a 1% w/v agarose gel. A total of 300 ng of each replicon sample was loaded onto the gel, and a voltage of 120 V was applied for a total runtime of 45 min. Gels were imaged using a ChemiDoc Imaging System (Bio-Rad, Hercules, CA, USA), and band intensity was quantified using Image Lab software version 6.1.0 (Bio-Rad). The percentage of the full-length replicon construct relative to a freshly prepared replicon control was estimated by normalizing the sample band intensity against the band intensity of the extracted replicon loading control on each gel. For whole replicon-NLC vaccine complexes, samples were also prepared without extraction via phenol:chloroform:isoamyl alcohol prior to treatment of samples with loading dye and heat. Evaluation of these whole complexes by agarose gel is an indicator of maintenance of the replicon nucleic acid binding the NLC, as the replicon that remains associated with NLC in the vaccine complex will not migrate into the gel and no distinct replicon band will be observed.

### NLC formulation lipid content analysis

NLC formulation lipid content was analyzed as described previously [21] with minor modifications. The concentrations of DOTAP, squalene, and trimyristin in the NLC were determined using a high-performance liquid chromatography (HPLC) method (1100 quaternary pump HPLC system, Agilent Technologies, Santa Clara, CA, USA) in combination with charged aerosol detection (Corona Veo Charged Aerosol Detector, Thermo Fisher Scientific). Two mobile phases were used for analysis: mobile phase A (60:15:25 methanol:chloroform:water, 10 mM ammonium acetate, and 1% w/v acetic acid) and mobile phase B (60:15:25 methanol:chloroform:isopropanol, 10 mM ammonium acetate, and 1% w/v acetic acid). Samples were diluted 1:10 with a solution of 50:50 methanol:chloroform. Subsequently, samples were further diluted 1:1 with mobile phase A to achieve a final 20-fold dilution. A reverse-phase HPLC method was used with an Atlantis T3 5 µm C18 100 Å column (Waters Corporation, Milford, MA, USA). The system was held at 35°C throughout the duration of the method, and the column was equilibrated with mobile phase A prior to sample injection. A 60 µL injection volume was used for each sample at a flow rate of 1 mL/min, and DOTAP, squalene, and trimyristin signal peaks were eluted using the gradient profile described in **Table 3**. Lipid concentrations were interpolated based on the peak area using a 5-point standard curve of each analyte and corrected for the dilution factor.

**Table 3.** Reverse-phase HPLC elution gradient profile.

| Time (min) | Channel A | Channel B |
| --- | --- | --- |
| 0 | 100% | 0% |
| 18.5 | 85% | 15% |
| 22.0 | 43.5% | 56.5% |
| 31.5 | 39.25% | 60.75% |
| 32.5 | 20% | 80% |
| 42.5 | 15% | 85% |
| 45.5 | 0% | 100% |
| 55.5 | 0% | 100% |
| 58.0 | 100% | 0% |
| 60.0 | 100% | 0% |

### *In vitro* cell uptake assay

To assess the replicon-delivery capacity of the spray-dried formulation, Vero cells (American Type Culture Collection, Manassas, VA, USA, #CCL-81) were transfected with green fluorescent protein (GFP)-expressing replicon complexed with NLC at an N:P of 10 (final nucleic acid concentration = 50 ng/µL) and spray dried using the same ultrasonic atomizer conditions described in the methods section above to generate a large powder. Vero cells were cultured in 6-well plates (35 mm well diameter) to approximately 80% confluence (approximately 1 million cells) the day of transfection. Vero cell monolayers were washed two times with warm phosphate-buffered saline (PBS). Following these washes, 500 µL of serum-free Dulbecco’s Modified Eagle Medium (DMEM) was added to each well and subsequently overlaid with 50 mg of spray dried powder containing the replicon-NLC complex. Upon addition to the cell-culture medium, the dry powder formulation rapidly dissolved, and the cells were incubated in this mixture at 37°C and 5% CO_2_ for 24 h. After the 24-hour incubation period, cells were overlaid with 2 mL DMEM + 10% fetal bovine serum, then imaged using an ImageXpress Pico imager (Molecular Devices, San Jose, CA, USA) to detect GFP expression post-transfection, using the 488 nm channel and an exposure time of 200 ms.

### *In vivo* immunogenicity

Mouse studies were performed under the oversight of the Bloodworks Northwest Research Institute’s (Seattle, WA, USA) Institutional Animal Care and Use Committee (IACUC), protocol #5389-01. All animal work followed applicable sections of the Final Rules of the Animal Welfare Act regulations (9 CFR Parts 1, 2, and 3) and the *Guide for the Care and Use of Laboratory Animals* [49]. Mice used in this study were female C57BL/6J, between 6-8 weeks of age, and purchased from The Jackson Laboratory (Harbor, ME, USA). All female mice were used to maximize the statistical power needed to detect differences in immunogenicity between vaccine formulations [50]. Immunization was performed via bilateral intramuscular injection into the rear quadriceps muscle (50 µL/leg, 100 µL total) at a total dose of 5 µg replicon for each sample tested. Blood was collected in microtubes (Sarstedt, Nümbrecht, Germany) by retro-orbital route immediately prior to immunization (Day 0) and 21 days post-immunization (Day 21) via terminal cardiac puncture and processed for serum isolation. H5-binding serum IgG antibodies were then assessed by enzyme-linked immunosorbent assay (ELISA) as previously described [22]. Plates were coated with 1 µg/mL recombinant H5 (Immune Technology Corp., New York, NY, USA, #IT-003-0051p) in PBS and incubated overnight at 4°C. An anti-HA A/Vietnam/1203/04 influenza virus (VN04-8) antibody (Rockland Immunochemicals, Limerick, PA, USA, #200-301-976) was used as a positive control.

## RESULTS

### Successful spray drying of replicon-NLC vaccine complexes requires the inclusion of L-leucine as a shell-forming excipient

Due to the relatively thermolabile nature of the replicon and the DOTAP lipid component of the NLCs, the H5 replicon-NLC vaccine complex was spray dried using relatively low temperature process conditions to mitigate any temperature-dependent degradation of the vaccine. To this end, a theoretical spray drying process model that takes into account dimensions of the spray dryer unit, formulation solids content, and drying gas fluid dynamics [41] was used to quickly identify process conditions that would result in a sufficiently dry product (< 4% moisture content) while using a low drying gas temperature of 65°C, which was previously demonstrated to be sufficient for successfully spray drying a tuberculosis vaccine candidate that was known to be susceptible to heat-induced degradation [44].

In addition to low-temperature processing, successful spray drying of the H5 replicon-NLC vaccine complex required the selection of a suitable amorphous glass-forming excipient to form a stable dry powder. Therefore, a vaccine complex solution containing 10% w/v trehalose (Batch #1) was first spray dried using the aforementioned process conditions, as trehalose is a well-known glass stabilizer in the spray drying of biologics [51–55]. However, spray drying of this formulation resulted in a highly adhesive powder that deposited in the processing equipment and subsequent poor powder recovery (**Table 4**). We next tested the use of alternative disaccharide excipients in the feedstock formulation, including 9% lactose (w/v) (Batch #2) and an 8%/2% trehalose/sucrose (w/v) mixture (Batch #3), to investigate whether an alternative glass stabilizer could improve dry powder recovery; however, similarly low powder recovery was obtained for each of these batches. As there was no apparent benefit to using alternative amorphous glass-formers, all subsequent formulations were tested using 10% (w/v) trehalose as the primary glass-forming excipient.

**Table 4.** Test feedstock formulation conditions and dry powder yields.

| Batch # | Replicon (ng/μL) | NLC<sup>b</sup> (w/v %) | N:P Ratio | Trehalose (w/v %) | Sucrose (w/v %) | Lactose (w/v %) | L-Leucine (w/v %) | Feedstock Volume (mL) | Feedstock pH | Dry Powder Yield |
| --- | --- | --- | --- | --- | --- | --- | --- | --- | --- | --- |
| 1 | 100 | 0.96 | 10 | 10 | 0 | 0 | 0 | 10 | 7 | <10% |
| 2 | 100 | 0.96 | 10 | 0 | 0 | 9 | 0 | 10 | 7 | <10% |
| 3 | 100 | 0.96 | 10 | 8 | 2 | 0 | 0 | 10 | 7 | <10% |
| 4 | 25 | 0.24 | 10 | 10 | 0 | 0 | 0 | 40 | 7 | <10% |
| 5<sup>a</sup> | 50 | 0.48 | 10 | 10 | 0 | 0 | 1.9 | 20 | 7 | 70% |
| 6 | 100 | 0.96 | 10 | 10 | 0 | 0 | 1.9 | 15 | 7 | 16% |
| 7<sup>a</sup> | 50 | 0.48 | 10 | 10 | 0 | 0 | 1.9 | 70 | 7 | 72% |

<sup>a</sup>Test batches #5 and #7 are the same formulation composition but prepared at different scales (by feedstock volume).
<sup>b</sup>NLC w/v % is the total percentage of the NLC components (solids and liquids) in the liquid feedstock formulation.

Next, to test the influence of total solids content on the recovered yield, the Batch #1 feedstock formulation was diluted 4-fold with formulation buffer containing 10% trehalose (w/v) prior to spray drying (Batch #4). Interestingly, although the apparent visual size of the collected dry powder particles was decreased compared with the more concentrated batches, the recovered powder yield remained less than 10%, indicating continued significant adhesion of the particles to the process equipment. Considering that under similar processing conditions, a much higher dry powder yield was observed for a spray dried squalene emulsion-based vaccine that used trehalose as the glass-forming excipient [56], the apparent visual decrease in powder size observed here, as well as the lack of improvement in the recovered powder yield following spray drying of a diluted liquid feedstock, suggested the vaccine complex may be inducing agglomeration of the dry powder. Thus, it was hypothesized that the inclusion of a suitable shell-forming excipient was necessary to prevent dry powder agglomeration.

To test whether inclusion of a shell-forming excipient could decrease agglomeration of the spray dried powder and improve powder recovery, L-leucine was added to the liquid feedstock formulation for use as a known shell-former commonly used in spray drying of biologics [57]. The original vaccine concentration was diluted 2-fold (50 ng/µL H5 replicon) to limit any potential cohesion and/or adhesion of the particles. In addition, along with maintaining trehalose as the main glass-forming excipient, L-leucine was included in the formulation as a shell-forming excipient at a near-saturating concentration of 19 mg/mL (1.9% w/v), which is necessary for this excipient to work well as a shell former (Batch #5) [58]. Spray drying of this new vaccine formulation resulted in the formation of a fine white powder and a substantially increased dry powder yield of 70%. This identified lead candidate formulation was then increased to the original nucleic acid concentration of 100 ng/µL H5 replicon (accompanied by necessary scaling of the total NLC content to maintain the N:P ratio) to determine if L-leucine was able to prevent significant loss of the material during the spray drying process at a higher vaccine concentration (Batch #6). However, the recovered powder yield decreased to 16%, suggesting an upper limit to the total amount of free (not in complex with replicon) NLC in the feedstock formulation before process yield becomes significantly impacted. Finally, in an initial test of the scalability of spray drying of the L-leucine-containing formulation, the liquid feedstock volume for the high-yield formulation was increased from 20 mL to 70 mL (Batch #7). This resulted in a dry powder yield of 72%, consistent with the observed dry powder yield for the smaller-scale process. Therefore, we selected this spray dried formulation for all subsequent studies (**Table 5**).

**Table 5.** Lead spray drying candidate formulation composition.

| Component | mg/mL | % w/w |
| --- | --- | --- |
| Tris | 0.10 | 0.08 |
| Trisodium Citrate | 1.38 | 1.10 |
| Trehalose | 100.00 | 79.40 |
| DOTAP | 1.00 | 0.79 |
| Sorbitan Monostearate | 1.23 | 0.98 |
| Trimyristin | 0.08 | 0.06 |
| Polysorbate 80 | 1.23 | 0.98 |
| Squalene | 1.25 | 0.99 |
| L-Leucine | 19.63 | 15.59 |
| Replicon | 0.050 | 0.04 |
| **Total Concentration** | **125.95** | **-** |

### The physical characteristics of the spray dried vaccine complex are suitable for pulmonary delivery

The lead spray dried formulation was next assessed for its morphological characteristics. The moisture content of the powder was determined to be 2.9% by Karl-Fischer titration, and dynamic vapor sorption analysis indicated that the powder would remain highly stable to crystallization at RH values less than 60% (**Figure S1**). L-leucine is expected to form a crystalline shell on the powder surface of spray dried biologics [59–62]. To assess whether this event occurred for the spray dried H5 replicon-NLC vaccine, Raman spectroscopy was used to assess the degree of L-leucine crystallization in the dry powder. We qualitatively observed sharp peaks characteristic of significant crystallization of L-leucine, while trehalose was maintained in an amorphous state (**Figure 1A**). Next, FESEM was used to qualitatively assess dry powder particle size and morphology. The dry powder particles had a distinct rugose surface, and the majority of the particles appeared to span 1 – 7 µm in diameter (**Figure 1B**), in excellent agreement with the median particle size determined by both laser diffraction and aerodynamic particle size measurements (4.28 µm) (**Figure S2**). At higher magnifications we observed the presence of small pores on the surface of the powder with diameters ranging from roughly 35 to 60 nm (**Figure 1C**). Closer inspection of the powder morphology revealed the formation of a distinct shell layer consistent with the anticipated crystallization of L-leucine on the powder surface. We also observed an asymmetrical interior containing pores of approximately the same diameter as the pores observed on the powder surface (**Figure 1D**).Taken together, these data indicate the successful formation of a dry powder with morphological characteristics advantageous for pulmonary delivery.

**Figure 1.**
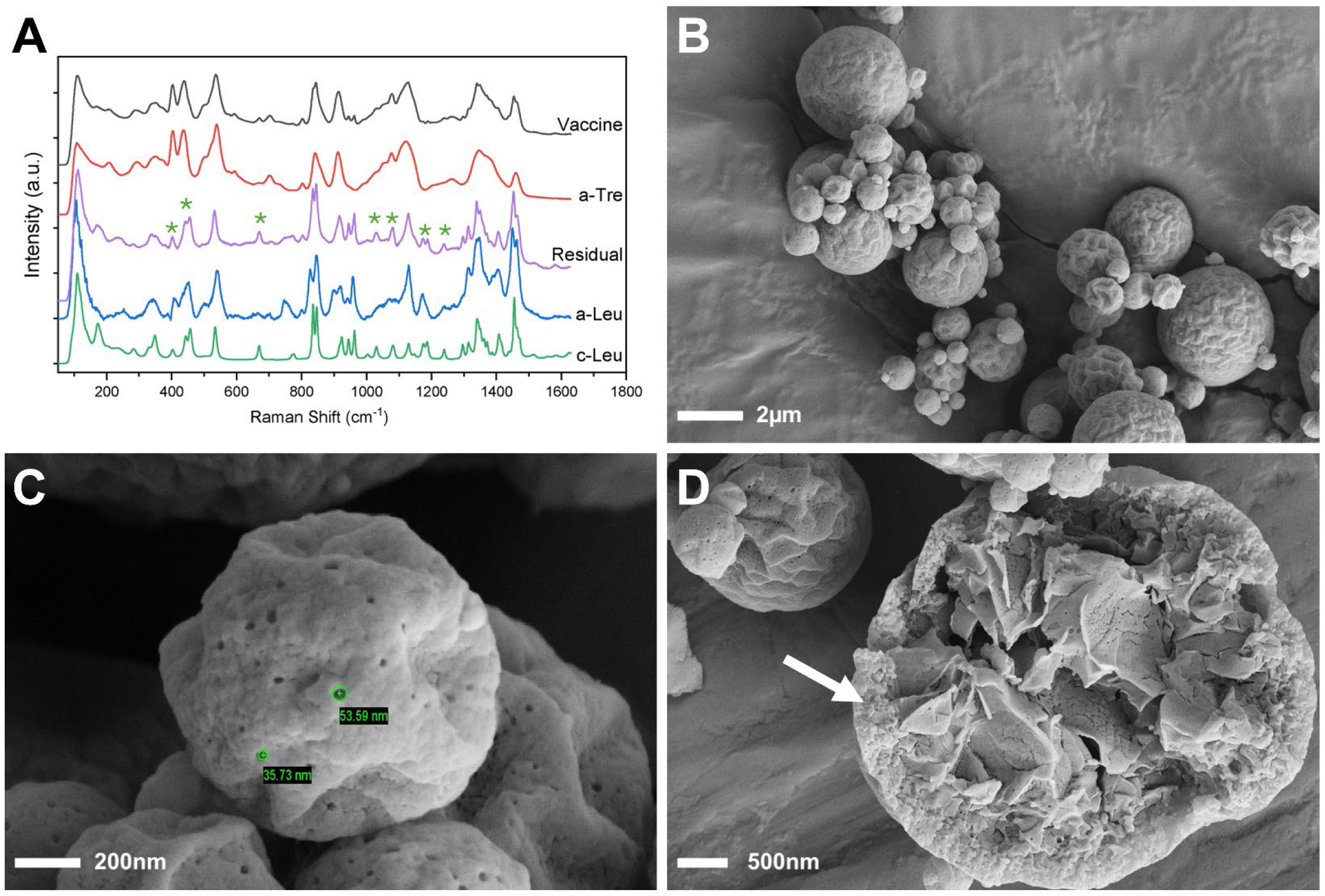
Characterization of spray dried replicon-NLC powder produced using a twin fluid atomizer. (A) Raman spectroscopy of the spray dried vaccine powder. The Raman spectrum for the complete spray dried vaccine (‘Vaccine’; black) was recorded, and the Raman spectrum for amorphous trehalose (‘a-Tre’; red) was subsequently subtracted from this spectrum to generate the residual spectrum (‘Residual’; purple). The residual spectrum was then compared with the raw Raman spectra for amorphous L-leucine (‘a-Leu’; blue) and crystalline L-leucine (‘c-Leu’; green). The green asterisks above the Residual spectrum denote peaks that are unique to the c-Leu Raman spectrum. (B) FESEM image of spray dried replicon-NLC vaccine complexes, indicating a generally spherical and rugose morphology. (C) FESEM image of a spray dried powder with green circles denoting example pores on the surface of the spray dried powder. Insets are approximate pore diameter in nm. (D) FESEM image of a particle that was intentionally ruptured for imaging of the powder interior. White arrow = solid shell that has formed on the powder surface.

### Spray dried vaccine complexes maintain suitable biophysical properties for *in vivo* immunogenicity

Next, the immunogenic and physicochemical properties of the spray dried vaccine were characterized after reconstitution of the dry powder to a replicon nucleic acid concentration of 50 ng/µL with nuclease-free water. The reconstituted vaccine was injected into C57BL/6 mice at a dose of 5 µg replicon, and H5-binding serum IgG antibody titers were assessed 21 days post-injection via ELISA. Antibody titers of the reconstituted vaccine were compared with those from control mice receiving a freshly prepared liquid (i.e., not spray dried) vaccine at the same replicon dose. Antibody titers of the spray dried and reconstituted vaccine were well maintained after the spray drying process, with levels only slightly below those detected for mice receiving the freshly prepared control vaccine (**Figure 2A**). Agarose gel densitometry indicated that the H5 replicon construct remained associated with the NLC lipids through the spray drying process (**Figure *2*B**). Organic extraction of the replicon from the spray dried complexes revealed a minimal apparent loss of full-length nucleic acid. An increase in the nanoparticle hydrodynamic diameter of the vaccine was observed after spray drying (**Figure 2C**) that is consistent with size increases that were previously observed for a lyophilized version of the replicon-NLC vaccine platform [21] and is likely the result of moderate aggregation of the lipid-based vaccine in the dried state. The apparent zeta potential of the spray dried vaccine showed a slight but acceptable increase of 2 mV relative to the freshly prepared liquid vaccine control (**Figure 2D**). Overall, after liquid reconstitution, the spray dried vaccine demonstrated immunogenicity and maintained acceptable physicochemical characteristics compared to a non-spray dried vaccine. We therefore decided to assess the long-term thermostability of the spray dried replicon-NLC vaccine powders to assess the degree of stability enhancement compared with the liquid vaccine.

**Figure 2.**
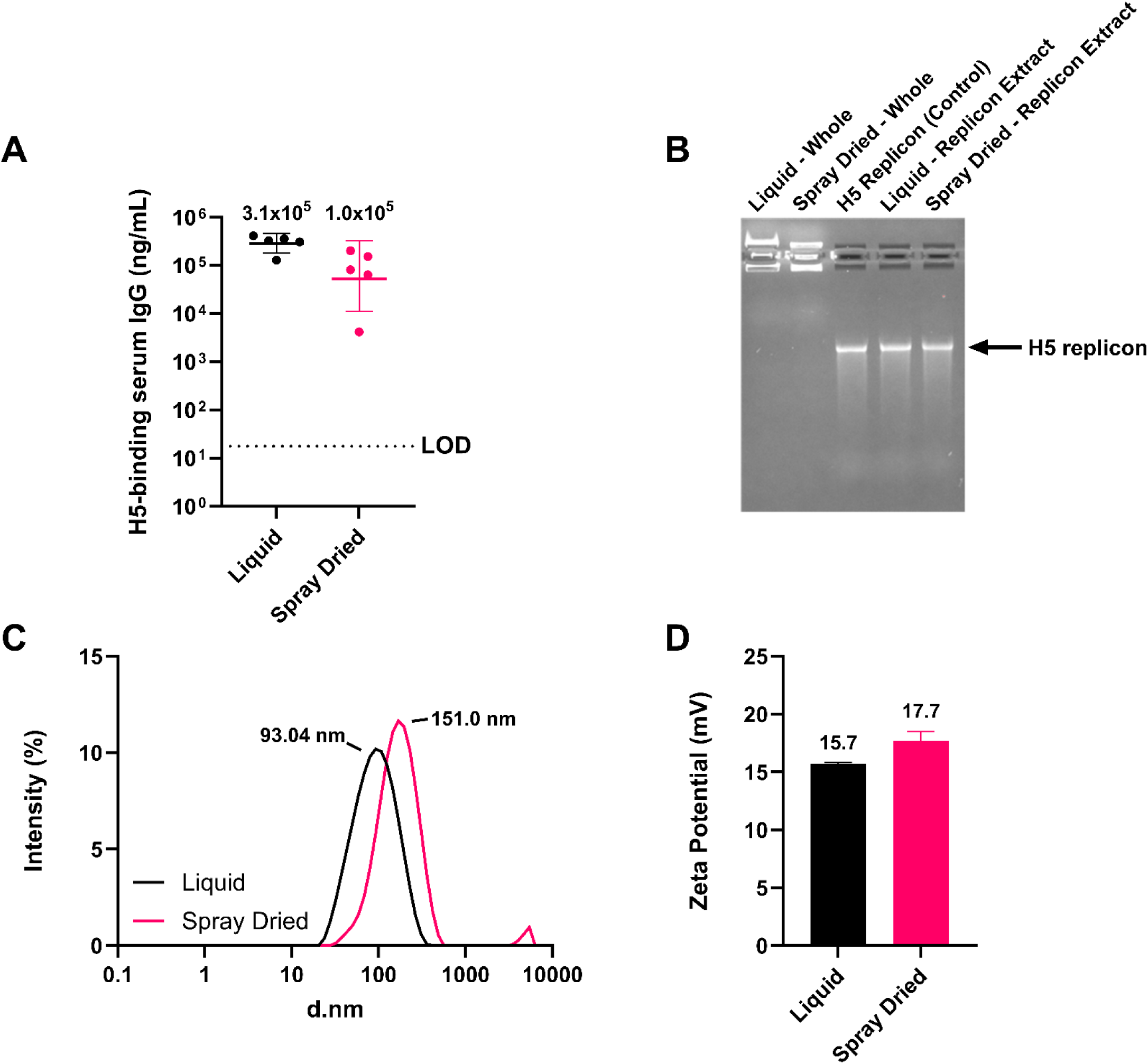
Biophysical and physicochemical characterization of spray dried and reconstituted replicon-NLC complexes, produced using a twin fluid atomizer, compared with a freshly prepared liquid vaccine (i.e., not spray dried). (A) Serum H5-binding IgG antibody titers from C57BL/6 mice, 21 days post-intramuscular dosing (5 µg replicon) with freshly prepared liquid vaccine or spray dried and reconstituted vaccine. Inset values correspond to the geometric mean measured antibody concentration. Data are presented as the geometric mean ± geometric standard deviation (*n* = 5 mice per group). LOD = limit of detection. (B) Agarose gel electrophoresis of freshly prepared liquid or spray dried and reconstituted replicon-NLC vaccine complexes. Liquid – Whole and Spray Dried – Whole lanes correspond to vaccine complexes that were not subjected to nucleic acid extraction from the NLC. H5 Replicon (Control), Liquid - Replicon Extract, and Spray Dried – Replicon Extract lanes correspond to vaccine complexes subjected to nucleic acid extraction. The black arrow denotes the expected size of the H5 replicon. (C) Dynamic light scattering intensity size distributions of freshly prepared liquid or spray dried and reconstituted vaccine complexes. Inset values correspond to the Z-average hydrodynamic nanoparticle diameter (*n* = 3 repeat measurements). (D) Zeta potential of freshly prepared liquid and spray dried-reconstituted vaccines. Inset values correspond to the mean measured zeta potential. Data are presented as the mean ± standard deviation (*n* = 5 repeat measurements).

### Spray dried powder morphology is well maintained after long-term storage at various temperatures

The morphology of the spray dried vaccine powder was assessed after storage for 3 months at 4, 25, and 40°C using SEM to determine the stability of the powder, which could have implications for long-term powder storage stability and dispersibility. Consistent with FESEM imaging (*vide supra*), at *t* = 0, the spray dried powder displayed an overall spherical morphology, with most of the particles appearing to be less than 10 µm in diameter (**Figure 3**). Dry powder morphology was well maintained after 3 months at all storage conditions tested, with no signs of powder fusing, indicating that the dry powder has high resistance to thermal-induced morphological changes.

**Figure 3.**
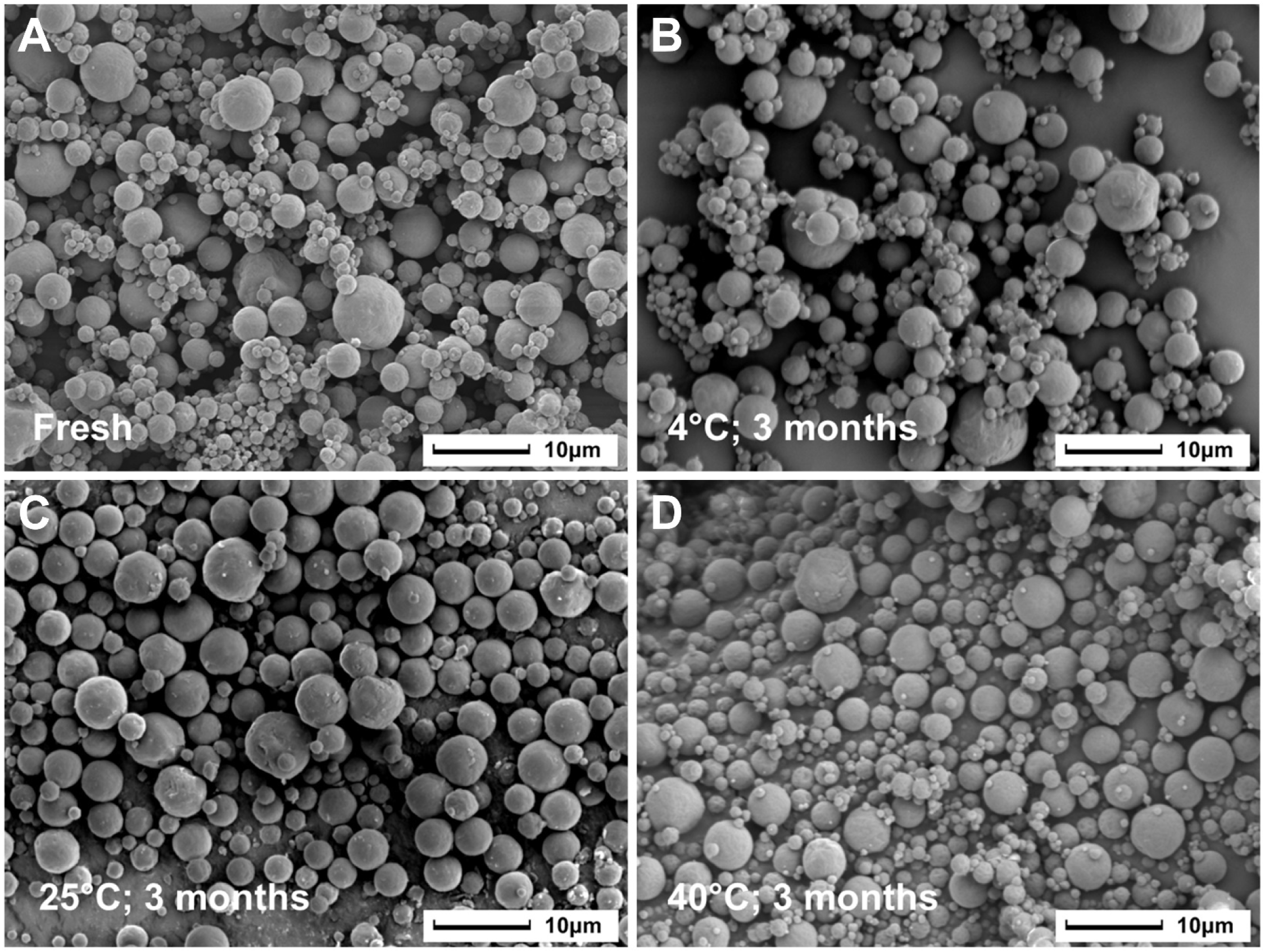
SEM imaging of dry powder vaccine morphology at various temperatures and timepoints. All images were captured at 2000x magnification and an acceleration voltage of 5 keV. (A) Spray dried powder morphology at *t* = 0, immediately after manufacturing. (B) Dry powder morphology after 3 months of storage at 4°C. (C) Dry powder morphology after 3 months of storage at 25°C. (D) Dry powder morphology after 3 months of storage at 40°C.

### Long-term physicochemical stability of the vaccine complex is well maintained upon spray drying

To assess the long-term physicochemical stability of the spray dried vaccine, the vaccine dry powder and the liquid feedstock vaccine control were both stored for 6 months at various temperatures, with monitoring of the physicochemical stability of the vaccine monitored throughout the duration of the study. At each timepoint, spray dried vaccine was reconstituted with nuclease-free water, as described above, prior to analysis. Full-length H5 replicon nucleic acid was rapidly degraded in the liquid vaccine control – the replicon showed complete degradation by agarose gel electrophoresis by 2 weeks, 1 month, and 3 months when stored at 40°C, 25°C, and 4°C, respectively (**Figure 1A**). By contrast, the spray dried vaccine showed substantially enhanced stability at all storage temperatures tested. Even after 6 months of storage at either 4°C or 25°C, full-length replicon nucleic acid was still detectable by agarose gel electrophoresis although at levels diminished from *t* = 0. Furthermore, the spray dried vaccine retained at least 80% of full-length nucleic acid after 1 month of storage at 40°C, and complete degradation was not observed until the 3-month time point even at this elevated storage temperature.

The chemical stability of the major NLC lipids in the spray dried vaccine powders (DOTAP, trimyristin, and squalene) was analyzed by reverse-phase HPLC after reconstitution of the dry powder to determine whether the spray drying process maintained the stability of the NLC vaccine component. All three major lipids showed excellent stability in the spray dried vaccine after 6 months of storage at 4°C and 25°C (**Figure 4B – D**, **Table S1**). When the dry powders were stored at 40°C, trimyristin continued to show excellent stability, whereas the more heat-labile DOTAP and squalene lipids began to show a gradual decrease in their total concentrations.

**Figure 4.**
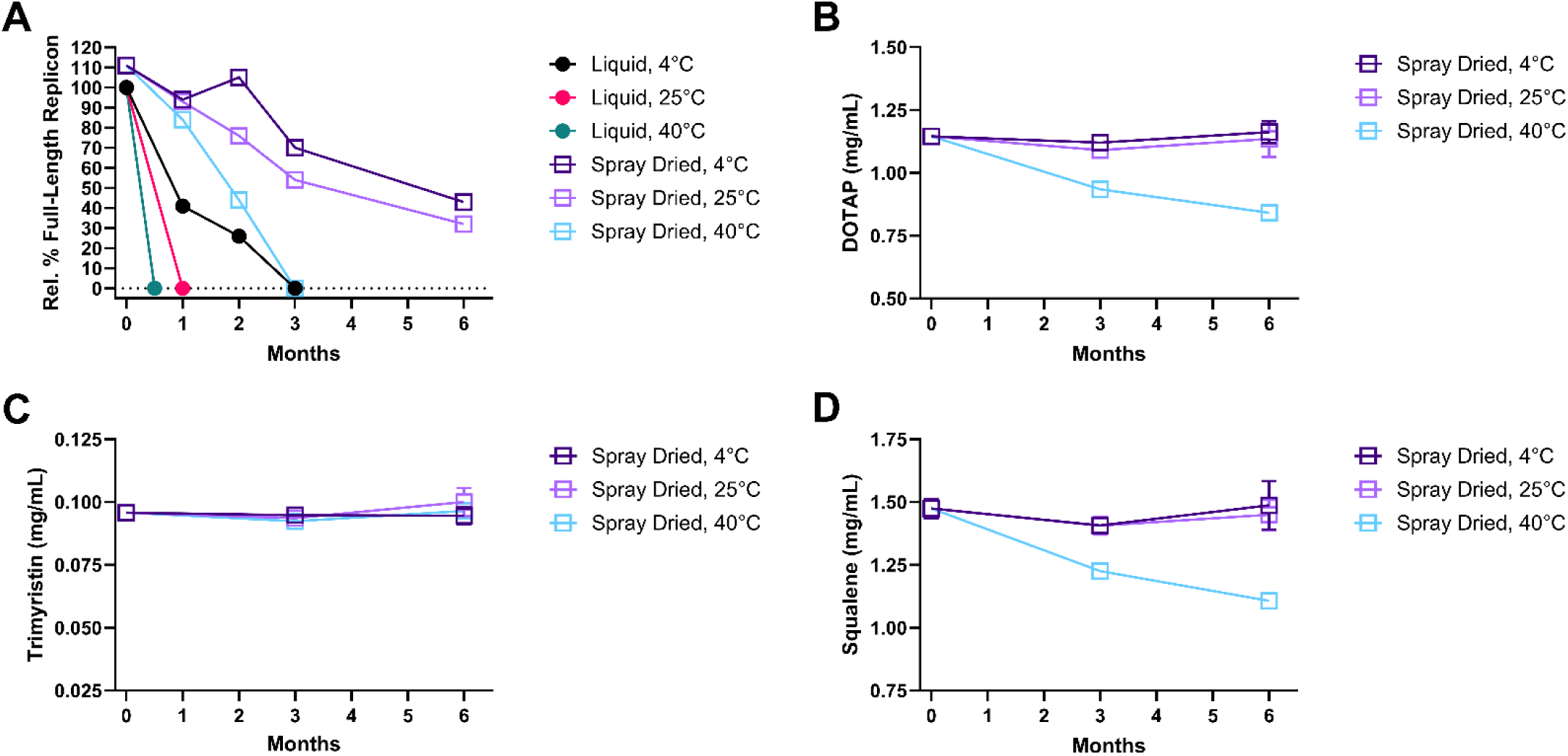
Chemical stability of the primary vaccine components after storage as a liquid or as a spray dried powder, followed by reconstitution immediately prior to analysis, at various temperatures and timepoints. (A) Percentage of the full-length replicon construct relative to a freshly prepared replicon control, as measured by agarose gel electrophoresis. (B) DOTAP content of spray dried vaccine complexes. (C) Trimyristin content of spray dried vaccine complexes. (D) Squalene content of spray dried vaccine complexes.

The hydrodynamic diameter, polydispersity index (PDI), zeta potential, and pH of the spray dried powder (after liquid reconstitution) and the liquid vaccine control were also assessed after long-term storage at various temperatures. Due to the increased thermolabile nature of the liquid vaccine relative to the spray dried vaccine, the liquid vaccine stored at the accelerated condition of 40°C was assessed more frequently and until complete replicon degradation was observed by agarose gel electrophoresis (**Figure 4A**). As noted previously, an initial size increase for the replicon-NLC vaccine complex in the spray dried vaccine was observed immediately after the spray drying process (**Figure 2C**). However, after this initial size increase at *t* = 0, the Z-average hydrodynamic diameter of the spray dried vaccine complex was well maintained during long-term storage (**Figure 5A**). Similarly, vaccine polydispersity was also well maintained for the spray dried vaccine at all monitored storage temperatures through 6 months (**Figure 5B**). By contrast, the liquid control showed a notable decrease in polydispersity after storage at 40°C for 2 weeks (**Figure 5B**). Nanoparticle zeta potential (**Figure 5C**) and sample pH (**Figure 5D**) were also well maintained for the dry powder vaccine at all storage temperatures monitored, indicating that the spray drying process did not adversely affect its physicochemical properties or long-term stability.

**Figure 5.**
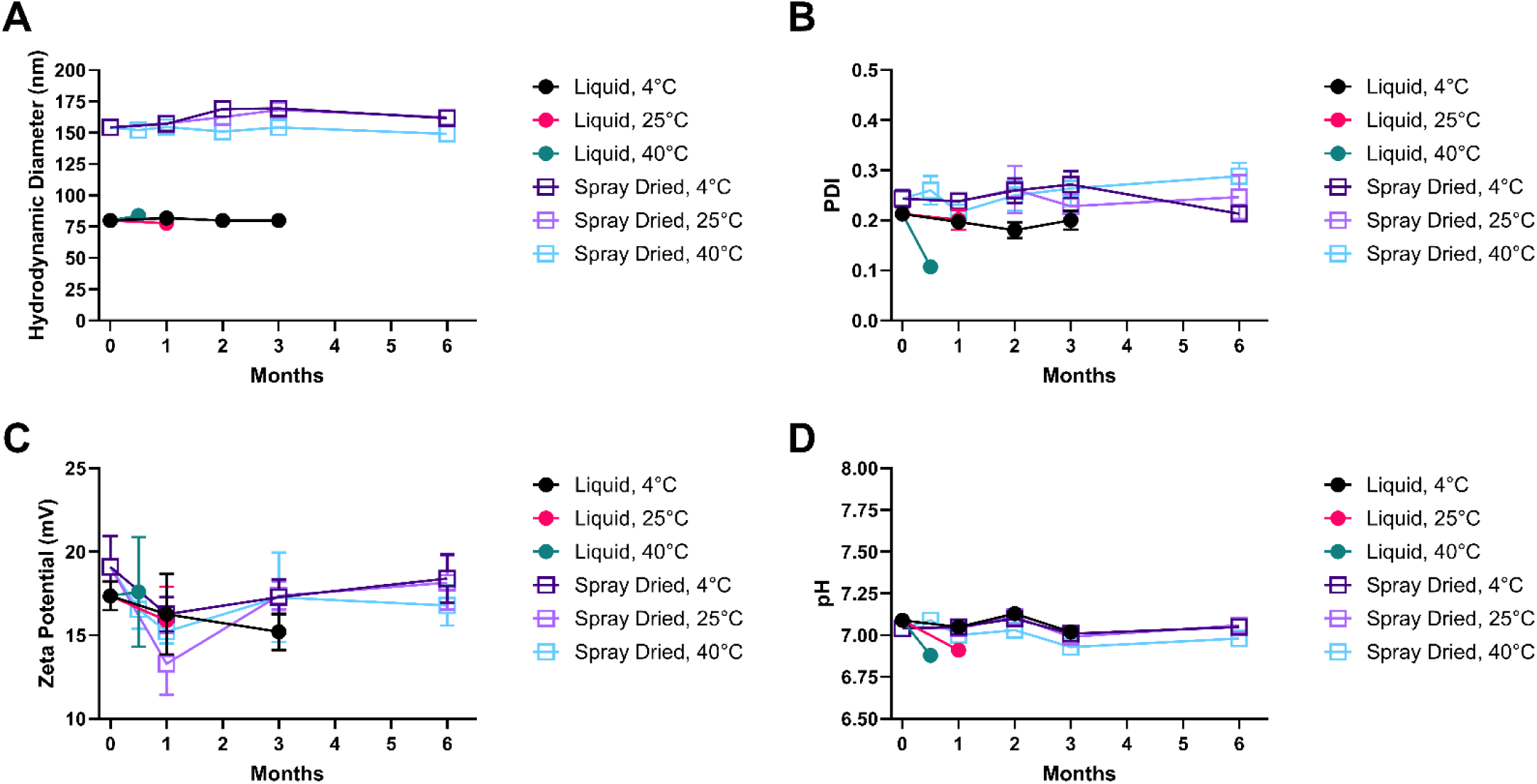
Physicochemical stability of spray dried-reconstituted or liquid H5 replicon-NLC vaccine complexes at various storage temperatures. (A) Hydrodynamic diameter of vaccine complexes as measured by DLS. (B) Polydispersity index of vaccine complexes. (C) Zeta potential of vaccine complexes. (D) pH stability of vaccine complexes.

### Spray dried replicon-NLC vaccine complexes remain immunogenic after several months of storage

We next assessed the long-term maintenance of immunogenicity of the spray dried vaccine. The spray dried H5 replicon-NLC vaccine was stored at 4°C under moisture-protective conditions. At various time points, the dry powder was reconstituted into a liquid and intramuscularly injected into C57BL/6 mice at a dose of 5 µg H5 replicon. H5-binding serum IgG antibody titers were measured 21 days-post injection. As a positive control, at each time point, fresh liquid vaccine was prepared and administered into mice via intramuscular injection, which showed consistent antibody titers with minimal spread throughout the duration of the study (**Figure S3**). Spray dried and reconstituted vaccine antibody titers were detectable at a level slightly lower than that of the fresh liquid vaccine preparation (**Figure 6**, **Figure S3**), consistent with previous observations (**Figure 2A**). Antibody titers were well maintained through 3 months of storage at 4°C, with waning immunogenicity not observed until the 6-month time point (**Figure 6**).

**Figure 6.**
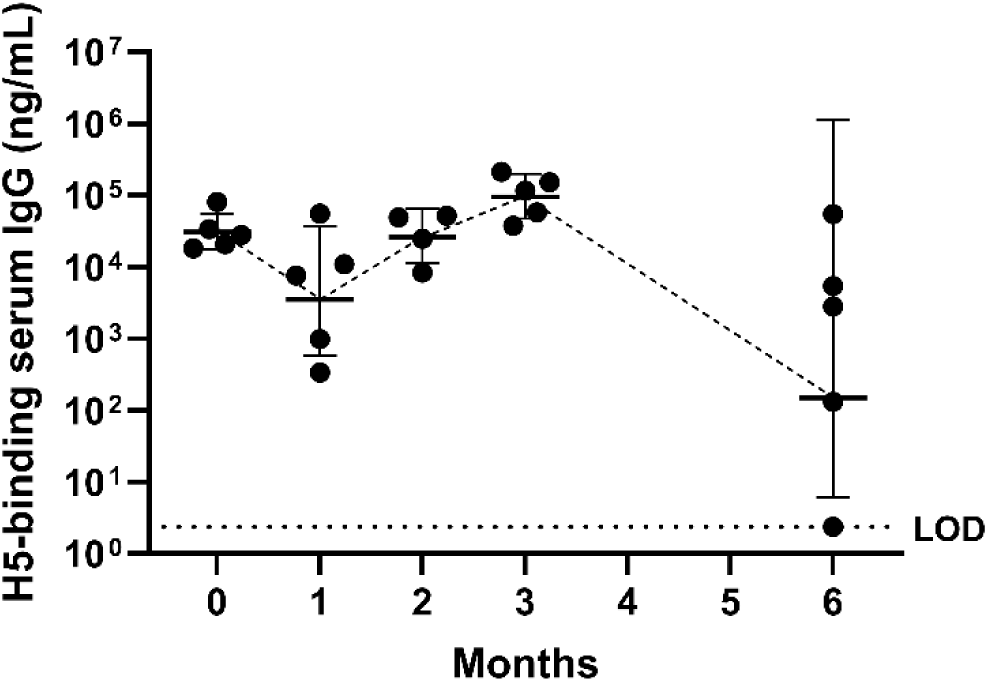
Immunogenicity of spray dried H5 replicon-NLC vaccine complexes produced using a twin fluid atomizer and stored at 4°C for 6 months. H5-binding serum IgG antibody titers in C57BL/6 mice receiving a 5 µg intramuscular dose of spray dried-reconstituted vaccine. Data are plotted as the geometric mean ± geometric standard deviation (*n* = 5 mice per group; a single statistical outlier at 2 months was identified using Grubbs statistical outlier test [α = 0.05] and removed prior to plotting of data). LOD = limit of detection.

### The size of the spray dried H5 replicon-NLC vaccine powder can be precisely controlled

A dry powder vaccine that is intended for exclusive delivery to the nose should have an MMAD greater than 20 µm – at this size, it is unlikely that a significant amount of the powder will reach the lungs upon intranasal administration, which could otherwise lead to potentially undesirable responses in the patient [44,63]. With a twin fluid atomizer, it is impossible for the spray drying system used here to produce a suitable dry powder that can meet the size criteria necessary for intranasal administration. We therefore spray dried the lead vaccine candidate formulation (**Table 5**) using a 25 kHz ultrasonic atomizer, which produces larger liquid droplets that will in turn result in a larger dry powder size. Spray drying of the vaccine using the ultrasonic atomizer resulted in a large initial droplet size (*d*_v,50_ = 62 µm, span = 0.81), ultimately producing a dry powder with an observed median volume equivalent diameter of 29 µm as assessed by SEM (**Figure 7A**) and laser diffraction analysis (**Figure S4**). Raman spectroscopy confirmed the presence of amorphous trehalose and crystalline L-leucine in the dry powder to a similar degree as the dry powder produced using the twin fluid atomizer, as anticipated (**Figure S5**). The physicochemical properties of this large powder were also consistent with those of the smaller vaccine powder produced using a twin fluid atomizer, including maintenance of full-length replicon nucleic acid through the spray drying process (**Figure S6**), an expected increase in the average hydrodynamic diameter of the vaccine nanoparticle upon spray drying (**Figure S7**), and a similar zeta potential to the liquid feedstock (**Figure S8**). Importantly, the large-sized vaccine powder elicited detectable and similar levels of H5-binding serum IgG antibody titers as the small-sized vaccine powder when administered via intramuscular injection to mice as a reconstituted liquid (**Figure 7B**), indicating that the dry powder size of the spray dried replicon-NLC vaccine platform can be adjusted over a wide range without compromising vaccine function.

**Figure 7.**
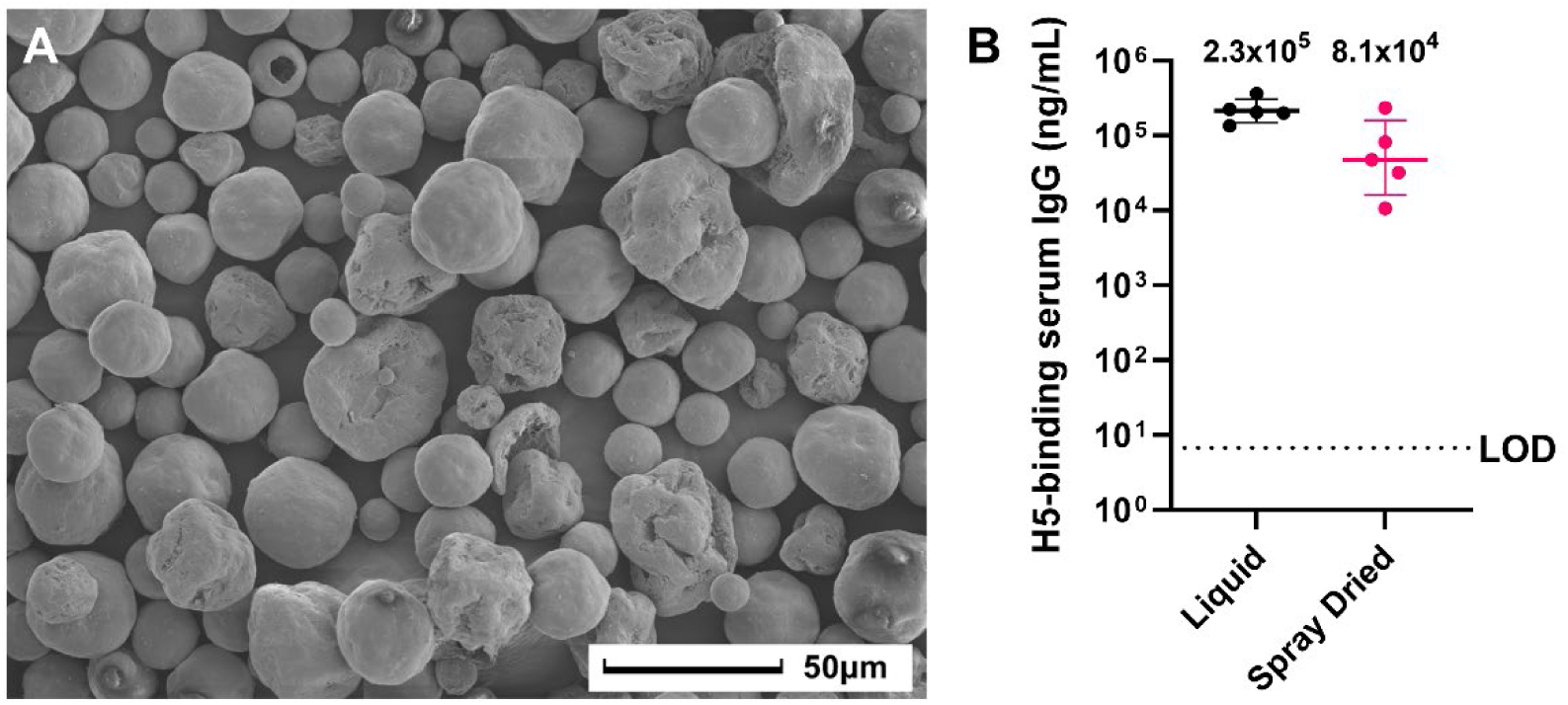
Characterization of spray dried H5 replicon-NLC vaccine complex produced using a 25 kHz ultrasonic atomizer. (A) SEM imaging of spray dried vaccine powder. (B) H5-binding serum IgG antibody titers from C57BL/6 mice, 21 days post-intramuscular dosing (5 µg replicon) with fresh liquid vaccine or spray dried-reconstituted vaccine. Inset values correspond to the geometric mean measured antibody concentration. Data are presented as the geometric mean ± geometric standard deviation (*n* = 5 mice per group). LOD = limit of detection.

To test whether spray dried replicon-NLC complexes can transfect cells when administered directly as a dry powder, replicon-NLC complexes were formed using a replicon construct that expresses GFP and subsequently spray dried using the ultrasonic atomizer to produce a large powder with similar morphological characteristics as the H5 replicon-NLC spray dried complex. After direct administration of 50 mg of the spray dried powder to Vero cells, fluorescence imaging revealed ubiquitous green fluorescence within the cells (**Figure S9**), confirming that cells are able to be transfected with a functional replicon construct *in vitro* when the spray dried complexes are administered as a dry powder without prior reconstitution. Taken together, these data highlight our ability to precisely tune the dry powder size of a functional replicon-NLC vaccine for a desired specific route of needle-free delivery (pulmonary or intranasal).

## DISCUSSION

Spray drying of the replicon-NLC platform represents a promising strategy for the rapid manufacturing and distribution of critical prophylactic vaccines to combat a global pandemic, as it would eliminate the need for strict cold-chain requirements for vaccine distribution. The spray dried vaccine would also help to alleviate vaccine hesitancy, as well as the strain placed on medical personnel, by obviating the need for needle-based administration. The present work demonstrates proof of concept that the replicon-NLC platform can be spray dried with precise control over the powder size while maintaining detectable vaccine immunogenicity. The physicochemical stability of the vaccine powder was well maintained over the course of several months owing to the addition of L-leucine, compared with the liquid vaccine control, which was quickly degraded at ambient temperatures. The spray dried vaccine also remained immunogenic in a mouse model during this same period, highlighting the potential for this spray dried platform to address the ongoing challenges with global vaccine distribution and uptake.

Upon spray drying of the replicon-NLC vaccine complex, the powder formed a crystalline L-leucine shell with an amorphous trehalose interior, and the biophysical properties of the nanoparticle after liquid reconstitution were found to overall be well maintained through the spray drying process, indicating that the process conditions selected for spray drying were adequate. The approaches described here demonstrate the utility of a theoretical and mechanistic understanding of spray drying thermodynamics and particle formation in choosing appropriate processing conditions. The use of a theoretical process model was critical in expediting process development for the spray drying of the replicon-NLC vaccine platform [41]. A mechanistic process model unique to a given spray dryer unit can be generated based on steady state energy and mass balances as well as empirical measurements of the heat loss throughout the dryer, enabling rapid technology transfer that is essential for robust pandemic response.

FESEM revealed the presence of small pores on the surface of the small spray dried replicon-NLC vaccine powder that was produced using a twin fluid atomizer. These pores had an apparent size ranging from roughly 35 to 60 nm, in good agreement with the expected size of the NLC vaccine component and suggesting the significant accumulation of the vaccine on the powder surface [20]. This could possibly explain the low process yield in the absence of L-leucine, as excessive accumulation of the replicon-NLC vaccine on the powder surface may promote powder agglomeration due to the surface-exposed lipids. L-leucine is an established crystalline excipient used to enhance the dispersibility of respirable powders, owing to its low solubility limit which promotes early crystal formation and subsequent rapid supersaturation at the droplet surface [61]. Friis and co-workers found that in order to increase processing yields of spray dried mRNA-LNPs, the inclusion of increasing amounts of another known dispersibility enhancer, trileucine, was essential in the formulation [33]. A similar phenomenon was observed by Sarode and co-workers, where process yields of a separate spray dried mRNA-LNP formulation were greatly enhanced by the inclusion of L-leucine [34]. In agreement with observations made by others, the inclusion of near-saturating amounts of L-leucine in the replicon-NLC formulations tested here substantially enhanced the dry powder yield, despite the presence of nano-sized pores on the powder surface observed by FESEM. This suggests that although L-leucine is not able to completely prevent the accumulation of vaccine nanoparticles on the droplet surface during drying, the early shell formation of L-leucine is likely able to limit the accumulation of the vaccine complexes on the droplet surface, resulting in lower surface enrichment of the vaccine and a final powder that is much less prone to agglomeration and subsequent processing loss [43]. Hence, the inclusion of a suitable shell-forming excipient at or near the solubility limit of the excipient appears to be required to achieve acceptable yields of spray dried replicon-NLC vaccine complexes (and other lipid-based nanoparticles), as these excipients compete with the apparently surface-active lipid-based nanoparticles during drying of the atomized feedstock liquid droplets.

In the context of pandemic response, it is critical that a spray dried vaccine maintain its morphological properties for prolonged periods of storage. If the dry powder particles begin to fuse or otherwise change morphology, this could have implications for the ability of the vaccine to be dispersed by the chosen administration device. Furthermore, chemical degradation of the replicon component of the vaccine due to undesirable moisture uptake (often indicated by fusing of powder particles) will ultimately result in decreased vaccine efficacy as the full-length antigen sequence is lost via hydrolysis. Here, upon long-term storage of the spray dried replicon-NLC vaccine complex, we observed that the vaccine powder morphology was well maintained for a minimum of 3 months at storage temperatures up to 40°C. This is likely due to the necessary inclusion of L-leucine in the feedstock formulation, which serves two purposes: (1) limiting the adhesiveness of the powder to improve process yield as described previously and (2) limiting the rate of moisture uptake to the powder through the formation of a protective crystalline L-leucine shell on the powder surface [57]. This further highlights the critical need of L-leucine in spray dried replicon-NLC formulations.

Another important consideration for spray dried vaccines in the context of pandemic response is the appropriate choice of dry powder particle size for the intended route of administration. Depending on the nature of the target pathogen, better disease outcomes may be achieved through either pulmonary or nasal routes of vaccine administration. To this end, we have demonstrated the ability to tune the anticipated aerodynamic particle size distribution of the spray dried H5 replicon-NLC vaccine to be suitable for either pulmonary or intranasal delivery, using either a twin fluid or ultrasonic atomizer, respectively. Upon liquid reconstitution and subsequent intramuscular administration, similar levels of H5-binding serum IgG antibody titers were elicited for both the small and large vaccine powder sizes, demonstrating the robustness of the spray drying process for the replicon-NLC vaccine platform. Spray dried vaccine immunogenicity was also well maintained for 3 months when the small vaccine powder was stored at 4°C, highlighting the long-term maintenance of vaccine functionality in a dry presentation. The measured antibody titers for both the small and large powders were slightly lower compared with titers generated using a freshly prepared liquid replicon-NLC vaccine control, which may be indicative of slight loss of the full-length nucleic acid to the spray drying process as hinted at by agarose gel electrophoresis. However, in the context of dry powder administration, this slight decrease in immunogenicity could be compensated for with further formulation and process development, or by simply adjusting the total administered powder dose downstream.

The ability to stably spray dry replicon-NLC vaccines with precise control over the dry powder size is an important and powerful development in the context of rapid pandemic response. While we were able to spray dry the replicon-NLC vaccine complex successfully by the inclusion of L-leucine, this study did not systematically evaluate process conditions and thus cannot conclude whether any changes to the process parameters used here would result in further improved powder yield or vaccine immunogenicity. Since it appears that the replicon-NLC vaccine complex readily collects on the droplet surface during the drying process, an increased understanding of the water surface activity of the vaccine platform will also be necessary to guide future choices of feedstock components and concentrations in an effort to maximize the dry powder yield without compromising vaccine efficacy, which will ultimately serve to further reduce costs associated with vaccine manufacturing. While the spray dried vaccine was able to maintain a detectable level of immunogenicity in a mouse model comparable to that of a liquid control, the present study only evaluated the vaccine via intramuscular administration as a reconstituted liquid. Ideally, a spray dried vaccine would be administered as a dry powder, obviating the need for tedious liquid reconstitution as well as needle-based delivery altogether. Testing of intranasal dry powder delivery in the future will require the use of a suitable animal model for dry powder vaccine administration, as well as an appropriate delivery device that is able to provide adequate moisture protection to the dry powder and suitable dispersion of the powder upon administration.

## CONCLUSIONS

We have successfully spray dried a replicon-NLC vaccine complex that remains immunogenic against H5 avian influenza. A shell-forming excipient such as L-leucine was successful in preventing the NLCs from collecting on the powder surface, thus improving overall yield. The spray dried complex demonstrates greatly enhanced thermostability over a liquid presentation, maintains good physicochemical stability, and remains immunogenic upon reconstitution and intramuscular administration in mice. Further, we can precisely control the size distribution of the spray dried powder, which potentially enables administration of the vaccine dry powder via pulmonary or intranasal routes, as desired. The ability to spray dry replicon-NLC vaccine complexes while maintaining replicon functionality has strong implications for the rapid manufacturing and global distribution of temperature-stable and efficacious vaccines in response to a global pandemic.

## ACKNOWLEDGEMENTS

We thank Jeffrey A. Guderian for their assistance with the manufacturing of the replicon constructs used for these studies. We also thank Dr. Valerie Soza for editing and proofreading the manuscript.

## AUTHOR CREDIT STATEMENT

**Wynton D. McClary**: Data Curation, Formal Analysis, Investigation, Methodology, Validation, Project Administration, Supervision, Validation, Visualization, Writing – Original Draft, Writing – Review & Editing; **John Z. Chen**: Formal Analysis, Investigation, Writing – Review & Editing; **Hui Wang:** Investigation, Formal Analysis, Writing – Review & Editing; **Ethan Lo:** Investigation, Writing – Review & Editing; **Julie Bakken:** Investigation, Writing – Review & Editing; **Joseph McCollum:** Investigation, Methodology, Writing – Review & Editing; **Christopher Press:** Resources, Writing – Review & Editing; **Eduard Melief:** Methodology, Resources, Writing – Review & Editing; **Devin S. Brandt:** Investigation, Writing – Review & Editing; **Andrew R. Martin:** Supervision, Writing – Review & Editing; **Reinhard Vehring:** Conceptualization, Methodology, Writing – Review & Editing; **Darshan N. Kasal:** Formal Analysis; Writing – Review & Editing; **Emily A. Voigt:** Conceptualization, Supervision, Writing – Review & Editing, Funding acquisition; **Alana Gerhardt:** Conceptualization, Supervision, Writing – Reviewing & Editing, Funding acquisition.

## DATA AVAILABILITY

Data will be made available on request.

