## Supplemental Information for "A Spray Dried Replicon Vaccine Platform for Pandemic Response"

**SUPPLEMENTARY MATERIAL**

**Table S1.** Spray dried vaccine lipid content after long-term storage at various temperatures

| | | **Lipid Content (mg/mL)** | | |
|---|---|---|---|---|
| **Component** | **Storage Temperature** | ***t* = 0** | **3 Months** | **6 Months** |
| DOTAP | 4°C | 1.15 | 1.12 | 1.16 |
| | 25°C | | 1.09 | 1.14 |
| | 40°C | | 0.93 | 0.84 |
| Trimyristin | 4°C | 0.10 | 0.09 | 0.09 |
| | 25°C | | 0.09 | 0.10 |
| | 40°C | | 0.09 | 0.10 |
| Squalene | 4°C | 1.47 | 1.41 | 1.49 |
| | 25°C | | 1.41 | 1.45 |
| | 40°C | | 1.23 | 1.11 |

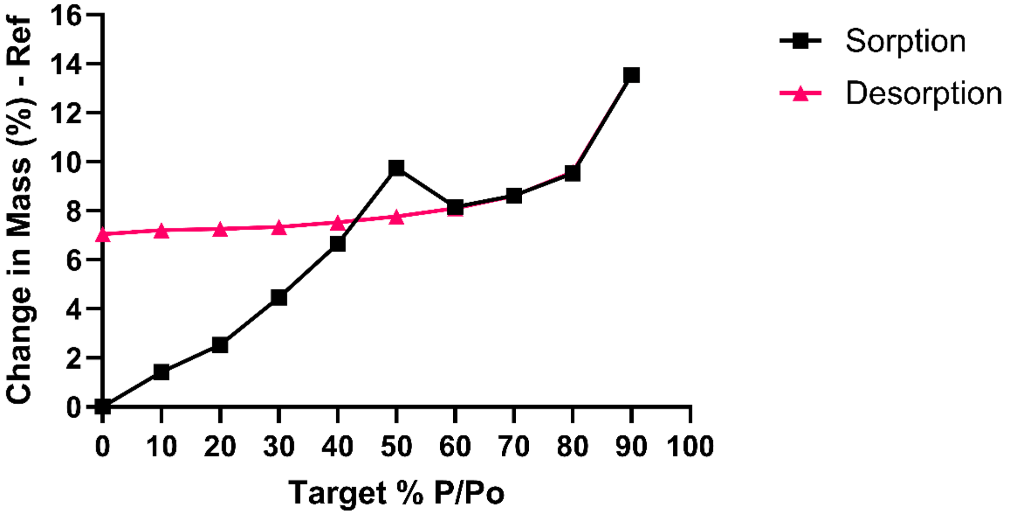


**Figure S1.** Moisture isotherm for spray dried replicon-NLC vaccine powder. Black trace – moisture sorption isotherm as the relative humidity is increased from 0% to 90%. Pink trace – moisture desorption isotherm as the relative humidity is decreased from 90% to 0%.

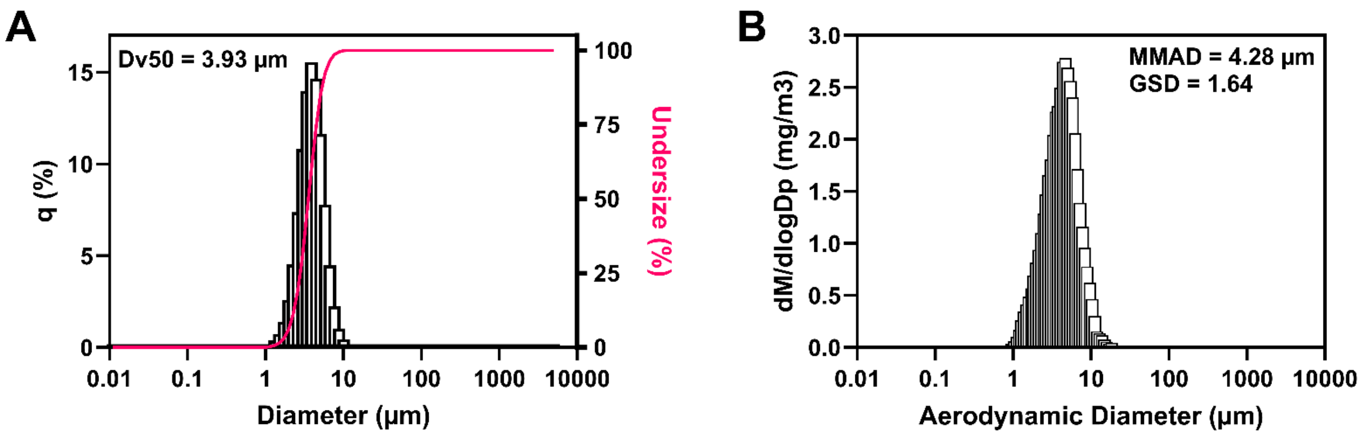


**Figure S2.** Particle size analysis of the spray dried powder produced using a twin fluid atomizer. (A) Laser diffraction measurement of the spray dried powder measured using 1-octanol as a non-solvent (Dv50 = median volume equivalent particle diameter). (B) Aerodynamic particle size measurement of the spray dried powder (MMAD = mass median aerodynamic diameter; GSD = geometric standard deviation).

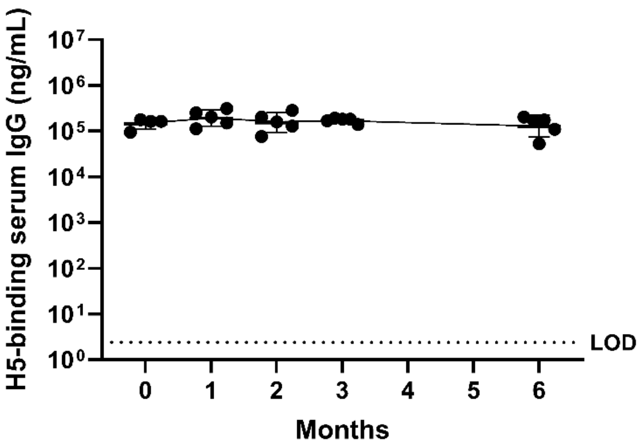


**Figure S3.** Immunogenicity of liquid H5 replicon-NLC vaccine controls prepared fresh on the day of each injection. H5-binding serum IgG antibody titers in C57BL/6 mice receiving a 5 µg intramuscular dose of spray dried-reconstituted vaccine. Data are plotted as the geometric mean ± geometric standard deviation (*n* = 5 mice per group; a single statistical outlier at 2 months was identified using Grubbs statistical outlier test [α = 0.05] and removed prior to plotting of data). LOD = limit of detection.

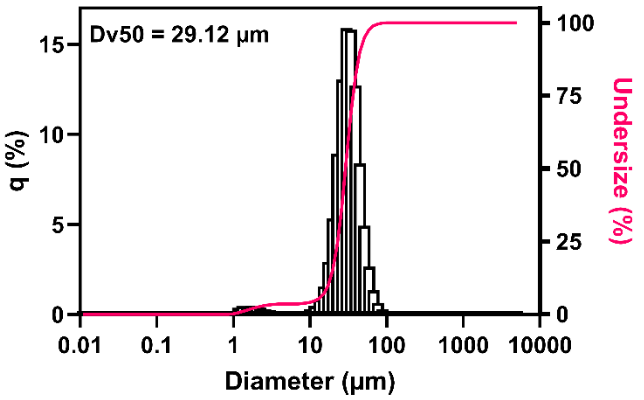


**Figure S4.** Particle size analysis of the spray dried powder produced using a 25 kHz ultrasonic atomizer, using laser diffraction analysis (Dv50 = median volume equivalent particle diameter).

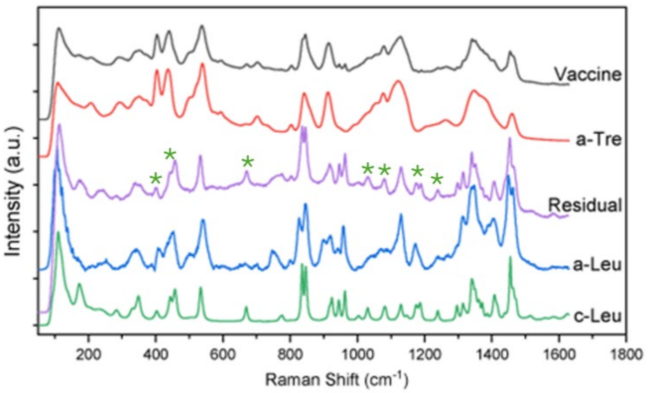


**Figure S5.** Raman spectroscopy of the spray dried vaccine powder produced using a 25 kHz ultrasonic atomizer. The Raman spectrum for the complete spray dried vaccine (‘Vaccine’; black) was recorded, and the Raman spectrum for amorphous trehalose (‘a-Tre’; red) was subsequently subtracted from this spectrum to generate the residual spectrum (‘Residual’; purple). The residual spectrum was then compared with the raw Raman spectra for amorphous L-leucine (‘a-Leu’; blue) and crystalline L-leucine (‘c-Leu’; green). The green asterisks above the Residual spectrum denote peaks that are unique to the c-Leu Raman spectrum.

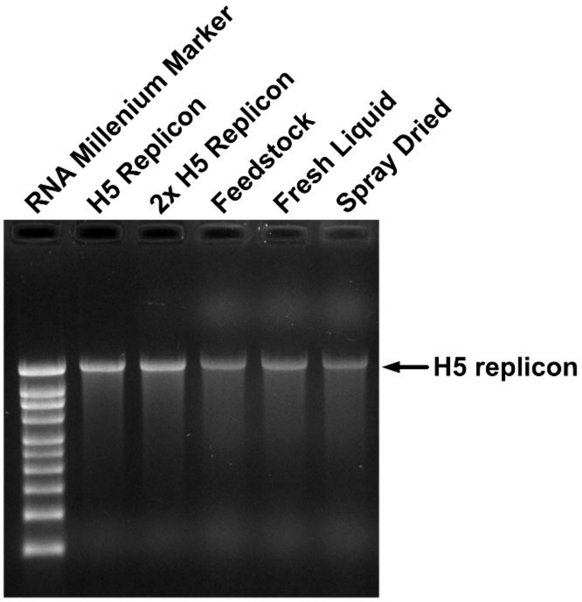


**Figure S6.** Agarose gel electrophoresis of liquid or spray dried and reconstituted replicon-NLC vaccine complexes produced using a 25 kHz ultrasonic atomizer. Samples were subjected to phenol:chloroform:isoamyl alcohol extraction prior to loading onto the agarose gel. H5 Replicon = freshly thawed replicon (Control); 2x H5 Replicon = nucleic acid at 2x the final concentration immediately prior to complexing with NLC; Feedstock = replicon from the feedstock solution immediately prior to spray drying; Fresh Liquid = nucleic acid extracted from a separately prepared replicon-NLC complex that was prepared fresh; Spray Dried = nucleic acid extracted from the spray dried and reconstituted dry powder. The black arrow denotes the expected size of the H5 replicon construct.

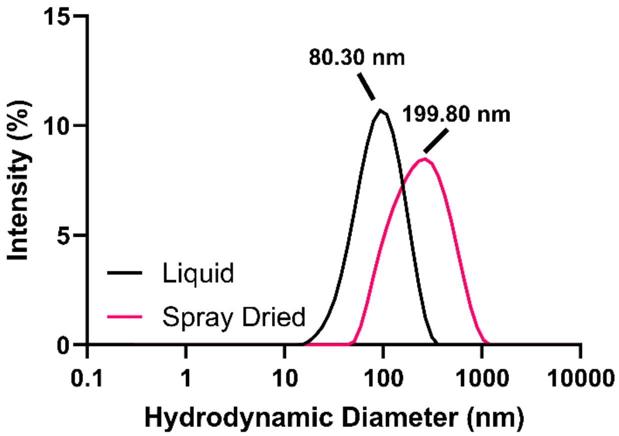


**Figure S7.** Intensity-based nanoparticle size distributions of freshly prepared liquid feedstock or spray dried and reconstituted replicon-NLC vaccine produced using a 25 kHz ultrasonic atomizer. Inset values are the Z-average diameter of the nanoparticles in solution. Data are shown as an average of $n = 3$ replicate measurements.

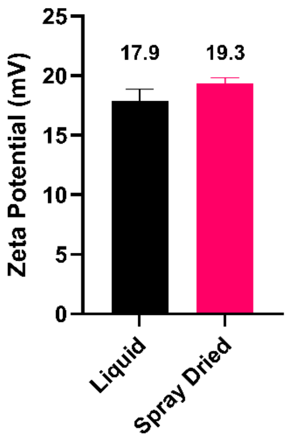


**Figure S8.** Zeta potential of freshly prepared liquid feedstock or spray dried and reconstituted replicon-NLC vaccine produced using a 25 kHz ultrasonic atomizer, as measured by electrophoretic light scattering. Inset values correspond to the mean measured zeta potential of liquid or spray dried and reconstituted vaccine complexes. Data are presented as the mean and standard deviation of $n = 5$ measurements.

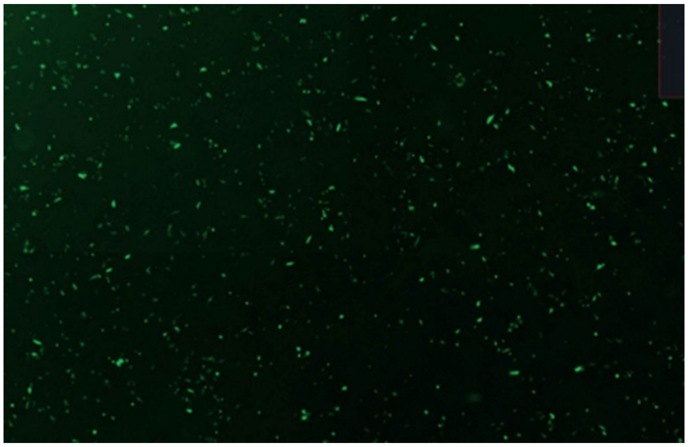

**Figure S9.** Fluorescence imaging of Vero cells after direct powder dosing for 24 h with 50 mg of a large spray dried powder containing replicon-NLC complexes that express green fluorescent protein. Expression is noted by the presence of green fluorescence within the cells.
